# Identification and Optimization of cell active 4-anilino-quin(az)oline Inhibitors for Protein Kinase Novel 3 (PKN3)

**DOI:** 10.1101/2020.03.02.972943

**Authors:** Christopher R. M. Asquith, Louisa Temme, Tuomo Laitinen, Julie Pickett, Frank E. Kwarcinski, Parvathi Sinha, Carrow I. Wells, Graham J. Tizzard, Reena Zutshi, David H. Drewry

## Abstract

The development of a small library of 4-anilinoquinolines led to the identification of 7-iodo-*N*-(3,4,5-trimethoxyphenyl)quinolin-4-amine **16** as a potent inhibitor of Protein Kinase Novel 3 (PKN3) with an IC_50_ of 1.3 μM in cells. Compound **16** presents a useful potential tool compound to study the biology of PKN3 including links to pancreatic and prostate cancer, along with T-cell acute lymphoblastic leukemia. These compounds may be useful tools to explore the therapeutic potential of PKN3 inhibition in prevention of a broad range of infectious and systemic diseases.

---

Protein kinase N (PKN, protein kinase novel) family genes encode for the three isoforms PKN1 (PKNα/PRK1/PAK1), PKN2 (PKNγ/PRK2/PAK2/) and PKN3 (PKNβ) that form the subfamily of AGC serine/threonine protein kinases.^[1]^ PKNs are closely related to novel isoforms of the protein kinase C family and are therefore also named PRKs (protein kinase C-related kinases).^[2,3]^ More in detail, the catalytic domain of the mammalian PKN is homologous to protein kinase C family members at its C-terminal region and contains three repeats of an antiparallel coiled-coil (ACC) domain and a C2-like domain at its *N*-terminal region.^[4,1]^ All these domains are well-conserved among the PKN isoforms but the enzymatic properties and tissue distributions of PKN1-3 differ.^[1]^ While PKN1 and PKN2 are expressed ubiquitously PKN3 is only found in restricted tissues including skeletal muscle, heart, liver and human cancer cell lines.^[2, 4-6]^

Full catalytic activation of PKN1-3 requires phosphorylation of the activation loop by phosphoinositide-dependent kinase 1 (PDK1) and turn motif phosphorylation.^[2, 4]^ PKN 1-3 differ in their responsiveness regarding activation by members of the small GTPase Rho, phospholipids, fatty acids and arachidonic acid.^[3, 4, 7]^ Mukai *et al*. investigated the physiological role of PKN3 in normal tissue with the help of PKN3 KO mice.^[7]^ It was shown that PKN3 is not required for development and growth to the adult stage, but lower migratory activity of embryonic fibroblast cells was observed suggesting PKN3 involvement in the actin cytoskeleton regulation of primary fibroblastic cells. ^[7]^ Additionally, normal vascular development does not require PKN3 but plays a role in angiogenesis support. Tumor angiogenesis is not inhibited by PKN3 KO.^[1]^

However, PKN3 is involved in other pathological processes. Thus, PKN3 has shown to interact with a mediator of tumor invasion and metastasis in epithelial cancers, RhoC,^[8]^ and with p130Cas, known to regulate cancer cell growth and invasiveness.^[9]^ PKN3 acts downstream of phosphoinositide 3-kinase (PI3K). Growth factor stimulation of normal cells leads to transiently activated PI3K, which is rapidly turned off by the phosphatase and tensin homolog (PTEN).^[5]^ However, most frequently PTEN is inactivated in human cancer resulting in overactivation of the PI3K pathway, which in turn upregulates PKN3 and increases metastatic behavior.^[9]^ This is one of the processes that mediate malignant cell growth in prostate cancer.^[5]^ Additionally, the malignant behavior of breast cancer cells *in vitro* increased by overexpression of exogenous PKN3.^[8]^ Since PKN3 can act downstream of PI3K signaling,^[5,8]^ therapeutics that selectively inhibit PKN3 could provide valuable insights in the role of PKN3 in cancer biology and represent a promising approach to target these cancer types that lack tumor suppressor PTEN function or rely on chronic activation of PI3K.^[5]^ Thus, PKN3 inhibition was observed to result in growth inhibition of prostate and breast cancer xenografts that were PI3K-driven.^[5,8, 10]^

An siRNA formulation (Atu027) that knocks down PKN3 *in vivo*, preventing liver and lung metastases in mouse models and inhibiting prostate and pancreatic cancer growth passed phase I clinical trial and has advanced to the ongoing phase I/II trial in advanced pancreatic cancer (NCT01808638).^[11, 12]^ Atu027 was well tolerated, no responses of the innate immune system, often a problem with siRNA formulations, were observed and the treatment was not restricted to a type of cancer. It was anticipated that beneficial effects could be seen in all vascularized metastatic cancers.^[12]^ However, dose limiting toxicity was not reached in the phase I clinical trial and the potential remaining drawbacks of this siRNA approach include enzymatic instability, off-target effects, challenges in tissue-specific access due to their negative charge and size as well as rapid liver clearance and renal excretion.^[12-15]^

Additionally this PI3K pathway was found to be overactivated in many types of leukemia,^[10, 16-18]^ the expression level of PKN3 was significantly increased in human T-cell acute lymphoblastic leukaemia (T-ALL) and PKN3 deletion slowed T-ALL development without affecting normal hematopoiesis.^[10]^ PKN3 was also found to be a possible regulator of neovascularization and its inhibition might be beneficial for vascular diseases like arthritis and age-related macular degeneration.^[7]^ Another possible therapeutic field covers bone diseases like rheumatoid arthritis and osteoporosis since, downstream of the Wnt5a-Ror2-Rho signaling pathway, PKN3 facilitates bone resorption.^[19]^ The development of a PKN3 chemical probe would significantly enhance the understanding around the role of PKN3.^[20]^ Despite some reported PKN3 inhibitors (Figure 1),^[7,21]^ wider structure activity relationships and kinome selectivity are yet to be explored.^[21-25]^

**Figure 1.**
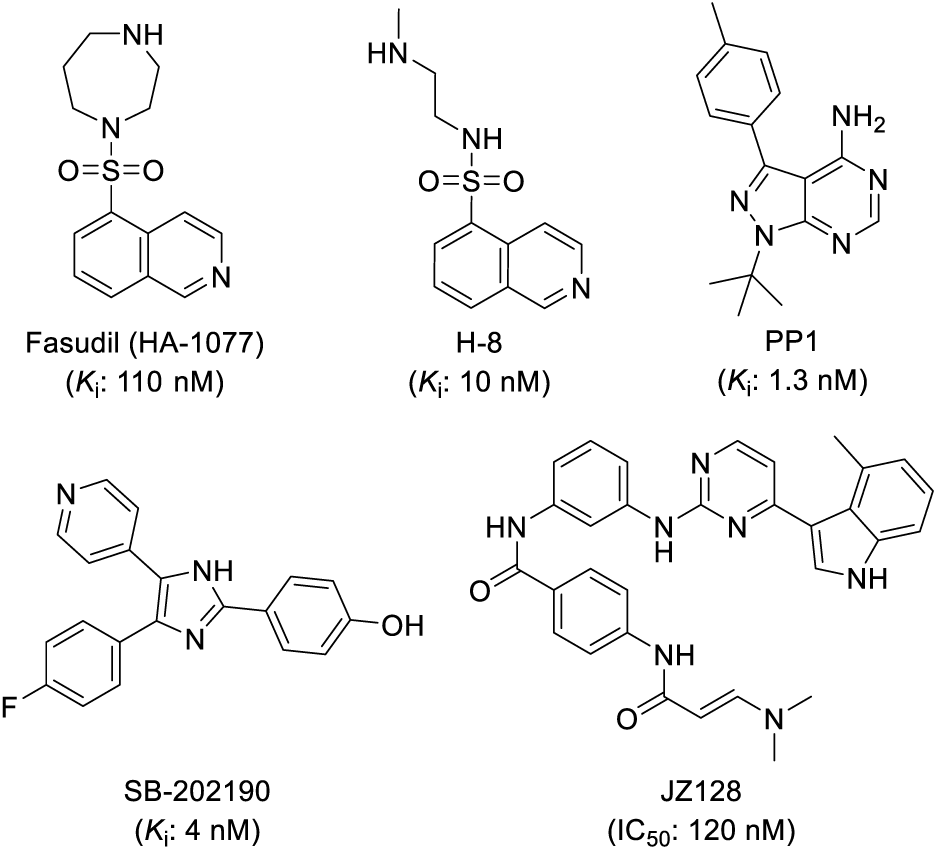
Previously reported compounds active against PKN3.

In order to develop a tractable lead compound for PKN3, we first looked at PKN3 inhibition using the well characterized second generation GlaxoSmithKline Published Kinase Inhibitor Sets (PKIS2).^[24-25]^ PKN3 was not part of the DiscoverX KinomeSCAN^®^ screen and is a historically understudied kinase, so we first screened PKIS2 against PKN3 in a split-luciferase binding assay^[29-30]^ and identified a set of 4-anilinoquinolines (**1**-**3**) with a common trimethoxyaniline motif (Figure 2). Compound **2** only hit four other kinases (GAK, RIPK2, ADCK3 and NLK) under 1 μM with PKN3 the third most potent (*K*_i_ = 280 nM). Compound **3** highlighted there was an ability to increase potency on PKN3 the signal digit nanomolar on PKN3. The kinome promiscuity profile of **3** (>18 kinases at 1 μM) was undesirable when looking for a narrow spectrum kinase inhibitor.

**Figure 2.**
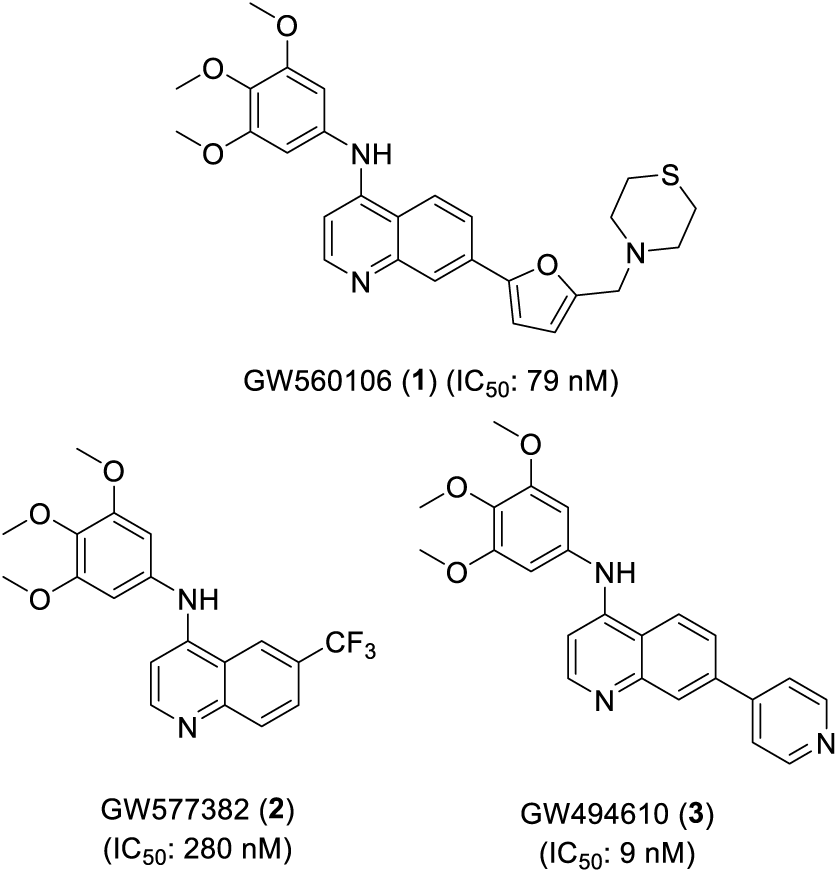
4-Anilinoquinolines identified as PKN3 inhibitors from PKIS2.

A small series of 4-anilinoquin(az)oline literature inhibitors was also screened in the PKN3 binding assay (Table 1). While cabozantinib and sapitinib were weak inhibitors of PKN3, gefitinib and lapatinib showed moderate activity. Vandetanib and tesevatinib revealed both IC_50_ values below 1 μM which further highlights the potential of this scaffold and the need for further structural modification to **2**, to improve its potency and selectivity towards PKN3.

**Table 1.** Literature 4-anilinoquin(az)oline PKN3 results.

| Compound | PKN3 $IC_{50}$ (μM)<sup>a</sup> |
| --- | --- |
| Cabozantinib | 42 |
| Sapitinib (AZD8931) | 44 |
| Gefitinib | 23 |
| Lapatinib | 5.9 |
| Vandetanib | 0.54 |
| Tesevatinib (XL-647) | 0.24 |

<sup>a</sup> $IC_{50}$ values generated in SLCA assay (n=2)<sup>[29-30]</sup>

To probe the structural requirements and improve potency for PKN3 inhibition we synthesized a series of analogs (Scheme 1). 4-Anilinoquinolines were prepared by heating the corresponding 4-chloroquinoline derivative and substituted aniline in ethanol to reflux overnight.^[31-35]^ The synthesis afforded products (**2, 4**-**57**) in good to excellent yield (24-93 %).

We first synthesized direct analogs of **2** with a two pronged strategy to improve potency on PKN3 and to overcome the innate GAK activity of this scaffold through divergent SAR.^[31-35]^ Analogs for GW577382 were previously used to generate a selective chemical probe for GAK.^[31-33]^ Hence, literature GAK nanoBRET data were used as a guide to GAK activity of the compounds as the GAK activity is the eventual probe quality benchmark for potency and selectivity. The compounds were initially screened for PKN3 binding in the split-luciferase binding assay (SLCA assay),^[29-30]^ in a 8-point dose response format to determine the IC_50_ values (Table 2).

**Scheme 1.**
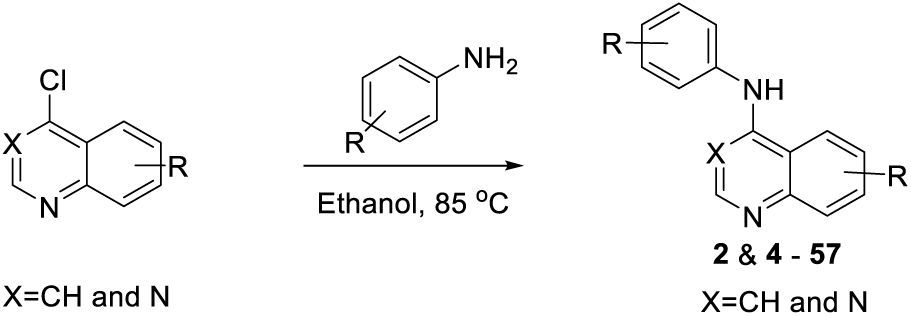
General synthetic route to analogs (**2** & **4** - **57**).

**Table 2.** Results of initial optimization of **2**.

| Cmpd | X | R<sup>1</sup> | R<sup>2</sup> | PKN3<sup>a</sup> | GAK<sup>b</sup> |
| --- | --- | --- | --- | --- | --- |
|  |  |  |  | IC<sub>50</sub> (μM) | in-cell IC<sub>50</sub> (μM) |
| 2 | H | CF<sub>3</sub> | H | 0.28 | 0.18 |
| 4 | N | CF<sub>3</sub> | H | 4.4 | 4.5 |
| 5 | H | H | H | 1.1 | 0.78 |
| 6 | H | F | H | 0.70 | 0.27 |
| 7 | N | F | H | 5.8 | 3.6 |
| 8 | H | Cl | H | 0.070 | 0.21 |
| 9 | H | Br | H | 0.0093 | 0.048 |
| 10 | H | I | H | 0.098 | 0.11 |
| 11 | N | I | H | 2.7 | 0.89 |
| 12 | H | H | F | 0.39 | 0.51 |
| 13 | N | H | F | 3.9 | >5 |
| 14 | H | H | Cl | 0.027 | 0.25 |
| 15 | H | H | Br | 0.12 | 0.27 |
| 16 | H | H | I | 0.014 | 0.074 |
| 17 | N | H | I | 0.93 | 2.4 |
| 18 | H | <sup>t</sup>Bu | H | 0.31 | 0.33 |
| 19 | H | CN | H | 0.21 | 0.35 |
| 20 | H | SO<sub>2</sub>CH<sub>3</sub> | H | 0.60 | 3.1 |
| 21 | H | OCH<sub>3</sub> | H | 0.74 | 0.28 |
| 22 | H | OCH<sub>3</sub> | OCH<sub>3</sub> | 0.48 | 0.025 |
| 23 | H | H | OCH<sub>3</sub> | 0.92 | 0.44 |
| 24 | H | H | CF<sub>3</sub> | 0.24 | 0.52 |
| 25 | N | H | CF<sub>3</sub> | 2.6 | >5 |
| 26 | H | H | CN | 0.079 | 0.34 |

<sup>a</sup>IC<sub>50</sub> values generated in SLCA assay (n=2)<sup>[26]</sup>; <sup>b</sup>in-cell IC<sub>50</sub> values obtained in NanoBRET assay and previously reported<sup>[31-33,35]</sup>

The synthesized 6-trifluoromethyl quinoline **2** was active on PKN3 (IC_50_ = 280 nM) and had potent GAK activity in cells (IC_50_ = 290 nM). A switch to quinazoline (**4**) resulted in a decrease of activity by 15-fold on PKN3, but 25-fold on GAK. Removal of the 6-trifluoromethyl moiety from the quinoline scaffold (**5**) also led to a 3-4-fold drop on both PKN3 and GAK compared to **2**.

However, addition of a fluoro group in 6-position (**6**) increased potency on GAK by 3-fold, with nearly no change on PKN3. The corresponding quinazoline **7** showed a similar decrease in activity on both kinases as with compound **4**. The addition of a chloro atom in 6-position (**8**) resulted in a 4-fold increased activity on PKN3 (IC_50_ = 70 nM) and a maintenance of GAK activity (IC_50_ = 210 nM). Increasing the size of the substituent to the bromo (**9**) realized a compound with single digit nanomolar activity (IC_50_ = 9.3 nM) and a 5-fold window over GAK in cells (IC_50_ = 48 nM) as well as a known narrow spectrum on the kinome.^[31-33]^ A further increase of the halogen atom size to the iodo (**10**) led to a decrease in activity compared to **9** in line with activities observed in the chloro-substituted compound **8**. The corresponding quinazoline **11** showed a drop off in PKN3 activity (27-fold) but a shallower drop off in GAK activity (8-fold).

The 7-fluoro isomer **12** showed a 2-fold lower potency than the corresponding 6-fluoro analog **6** on GAK. The quinazoline analog **13** revealed a sustainable lower IC_50_ value which is consistent with the potency of previous analogs **4, 7** and **11**. The increase of the atom size to a chloro atom in this 6-position of **14** resulted in the first compound that showed an increased selectivity between PKN3 and GAK with a 10-fold difference in activities (IC_50_ = 27 nM vs 250 nM). Surprisingly, the bromo derivative **15** reduced the selectivity to 2-fold, driven by a 4-fold decrease in PKN3 and a maintaining of GAK activity. The iodo analog **16** was potent on PKN3 (IC_50_ = 14 nM) but only 5-fold selective over GAK in cells. The quinazoline analog **17** showed a corresponding drop off in activity, but still showed PKN3 activity under 1 μM.

The 6-position *tert*-butyl isomer **18** revealed equipotent activity compared to **2** with no overall improvement. This was the same in case of the cyano analog **19**. However, 6-methyl sulfone-substituted compound **20** showed 5-fold selectivity towards GAK and moderate potency against PKN3 (IC_50_: 600 nM).

The 6-methoxy (**21**), 6,7-dimethoxy (**22**) and 7-methoxy (**23**) substituted quinolines showed a preference for GAK which is consistent with previously observed SAR showing high potency on GAK.^[29]^ A switch to the 7-trifluoromethyl moiety (**24**) was in line with the activities observed for PKN3 and GAK showing a 3-fold reduction in GAK activity compared to **2**. The quinazoline analog **25** showed a consistent sharp drop off in activity for both kinases. The 7-cyano analog **26** was potent on PKN3 (IC_50_ = 79 nM) with 4-fold selectivity over GAK.

In order to find potent analogs that lack GAK activity, we decided to take a look at a small library of lapatinib derivatives following up from the screening of compounds in table 1 and 2. Therefore, a simplified lapatinib scaffold with a 6,7-dimethoxyquin(az)oline motif (**27**-**34**) was used (Table 3). We know that GAK activity would be removed with addition of substitutions beyond smaller simple groups at the *para*-position of the aniline.

**Table 3.** Simplified lapatinib analogs **27**-**34**.

| Cmpd | X | R<sup>1</sup> | R<sup>2</sup> | R<sup>3</sup> | R<sup>4</sup> | PKN3<sup>a</sup> | EGFR |
| --- | --- | --- | --- | --- | --- | --- | --- |
|  |  |  |  |  |  | IC<sub>50</sub> (μM) | in-cell IC<sub>50</sub> (μM) |
| Lapatinib | CH | OCH<sub>3</sub> | OCH<sub>3</sub> | Cl | A | 5.9 | 0.021<sup>b</sup> |
| **27** | CH | OCH<sub>3</sub> | OCH<sub>3</sub> | Cl | A | 8.2 | 1.7<sup>b</sup> |
| **28** | N | OCH<sub>3</sub> | OCH<sub>3</sub> | Cl | A | 17 | 0.015<sup>b</sup> |
| **29** | CH | OCH<sub>3</sub> | OCH<sub>3</sub> | OH | B | 38 | 14<sup>c</sup> |
| **30** | CH | OCH<sub>3</sub> | OCH<sub>3</sub> | OH | C | >50 | 13<sup>c</sup> |
| **31** | CH | OCH<sub>3</sub> | OCH<sub>3</sub> | H | D | >50 | >20<sup>c</sup> |
| **32** | CH | OCH<sub>3</sub> | OCH<sub>3</sub> | H | E | >50 | >20<sup>c</sup> |
| **33** | CH | OCH<sub>3</sub> | H | OH | B | >50 | 14<sup>c</sup> |
| **34** | CH | H | OCH<sub>3</sub> | OH | B | 11 | 6.3<sup>c</sup> |

<sup>a</sup>IC<sub>50</sub> values generated in SLCA assay (n=2)<sup>[29-30]</sup>; <sup>b</sup>in-cell assay as previously reported<sup>[33]</sup>; <sup>c</sup>in-cell assay as Previously reported<sup>[36]</sup>

Switching from lapatinib to the simplified quinoline derivative **27** maintained activity on PKN3 and reduced in-cell EGFR activity by 80-fold.^[33,36]^ The quinazoline derivative **28** saw a 2-fold drop in PKN3 activity and an 80-fold boost in EGFR activity as previously reported. A few changes to the compound to expand the chemical space led to a further reduction in activity (**29**-**31**). The acetamide derivative **32** demonstrated that the benzyl group is important for activity since **32** showed no binding on either PKN3 or EGFR.^[36]^ Interestingly, the removal of the 7-position methoxy group of **29** resulting in analog **33** revealing no PKN3 inhibition improvement; however removal of 6-position methoxy in case of **34** showed a boost of activity of at least 5-fold (IC_50_ = 11 μM). Thus, the PKN3 activity is equipotent with in-cell EGFR activity (IC_50_ = 6.3 μM).^[36]^

We then considered that the halogen atom in the *meta*-position may have an effect on the activity profile of the scaffold. In the resulting series GAK is the collateral target of interest.^[31-35]^ A small series of halogenated analogs (**35**-**39**) was synthesized to probe this rational (Table 4). Replacing the methoxy groups of **2** and **9** with fluoro atoms resulted in analogs **35** and **36** with occupied *meta*-positions; both quinolines showed limited activity on both PKN3 and GAK.^[35]^ Removing the substitution of one of the two meta-positions resulted in quinolines **37** and **38** with limited activity on both kinases.^[35]^ However, switching back to the trimethoxy substitution and moving the fluoro atom to the 5- and 7-position yielded quinoline **39** with 8-fold selectivity towards GAK over PKN3 and potency below 1 μM.^[35]^

**Table 4.**
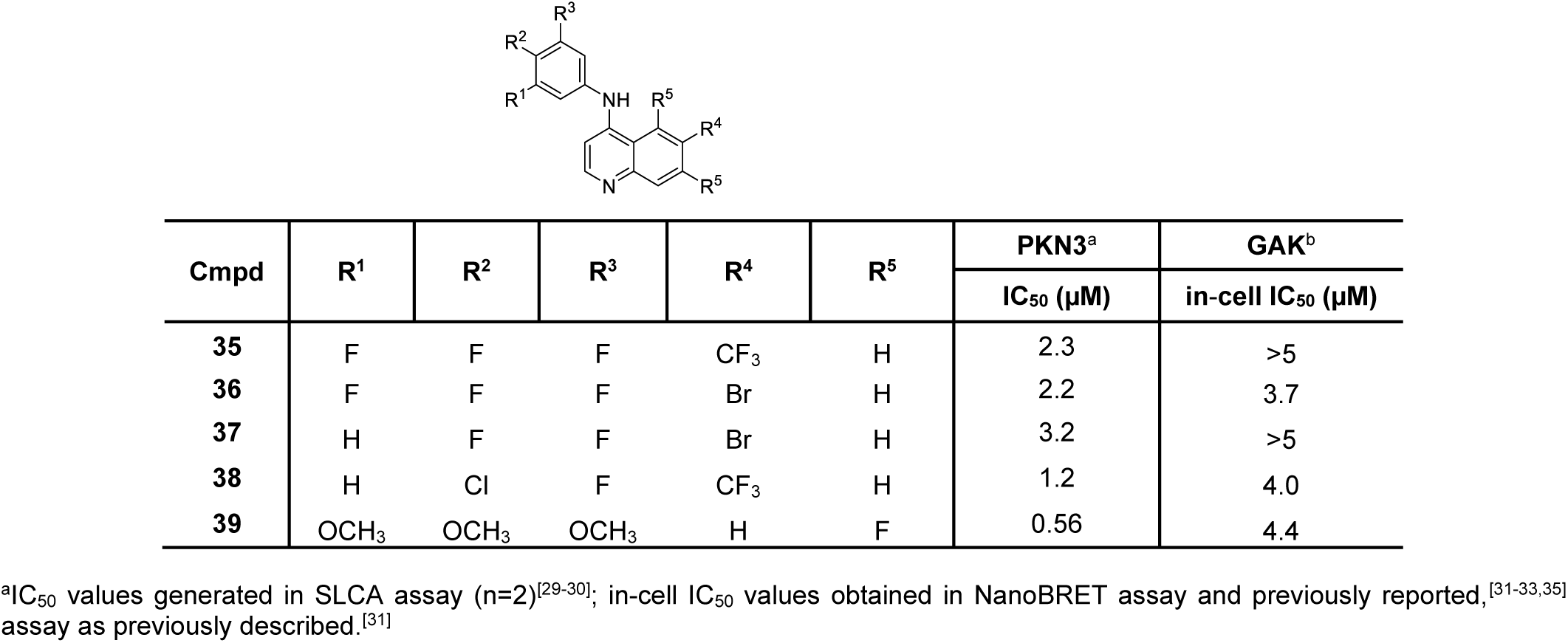
Results of halogenated quinolines **35**-**39**.

In an attempt to remove the GAK activity from the scaffold while maintaining PKN3 activity the addition of steric bulk in *para*-position of the preferred 6-bromo, 6,7-dimethoxy and 5,7-difluoro quinolines was investigated. These compounds (**40**-**57**) demonstrated only limited activity on PKN3 in an initial 1 μM (n=2) signal measurement (Table 5). The smaller *para*-substitutions including ethoxy (**40**-**42**), *iso*-propyl (**43**-**45**) and cyclobutyl (**46**-**48**) were all inactive at 1 μM. The *para*-benzyl substituted quinolines **49**-**51** showed a hint of activity but not in case of 5,7-difluoro derivative **51**. A further investigation of the 6-bromo and 6,7-dimethoxy quinolines and altering the electronics of the benzyl ring system. This yielded analogs (**52**-**57**) with activities similar to lapatinib, but with a higher ligand efficiency due to the removal of the solvent exposed region of lapatinib.

**Table 5.** Investigation of matched pairs of *para*-substituted quinolines.

| Cmpd | R<sup>1</sup> | R<sup>2</sup> | R<sup>3</sup> | R<sup>4</sup> | PKN3<sup>a</sup> |
| --- | --- | --- | --- | --- | --- |
|  |  |  |  |  | % control remaining at 1 μM |
| Lapatinib | OCH<sub>3</sub> | OCH<sub>3</sub> | H | - | IC<sub>50</sub> = 5.9 μM |
| **40** | OCH<sub>3</sub> | OCH<sub>3</sub> | H |  | 100 |
| **41** | Br | H | H |  | 99 |
| **42** | H | F | F |  | 100 |
| **43** | OCH<sub>3</sub> | OCH<sub>3</sub> | H |  | 97 |
| **44** | Br | H | H |  | 100 |
| **45** | H | F | F |  | 100 |
| **46** | OCH<sub>3</sub> | OCH<sub>3</sub> | H |  | 93 |
| **47** | Br | H | H |  | 100 |
| **48** | H | F | F |  | 100 |
| **49** | OCH<sub>3</sub> | OCH<sub>3</sub> | H |  | 92 |
| **50** | Br | H | H |  | 93 |
| **51** | H | F | F |  | 100 |
| **52** | OCH<sub>3</sub> | OCH<sub>3</sub> | H |  | 73 |
| **53** | Br | H | H |  | 100 |
| **54** | OCH<sub>3</sub> | OCH<sub>3</sub> | H |  | 93 |
| **55** | Br | H | H |  | 91 |
| **56** | OCH<sub>3</sub> | OCH<sub>3</sub> | H |  | 79 |
| **57** | Br | H | H |  | 95 |

<sup>a</sup> SLCA Assay (n=2)<sup>[29-30]</sup>**Table 6.** PKN3 NanoBRET results of the most potent analogs.

The compounds with the lowest IC_50_ value in the SLCA assay (**8**-**10** & **14**-**16**) were screened in a PKN3 NanoBRET assay to test their in-cell target engagement (Table 6).^[27-28]^ The NanoBRET consists of a Nano Luciferase, an extremely bright and small luciferase, fused to the *N*-terminal of PKN3 and transiently expressed in HEK293 cells. A red-shifted dye tethered to a fragment shown to bind the ATP binding site of PKN3 serves as the cell permeable tracer. This dye added together with the Nano Luciferase enzyme substrate produces a bioluminescence resonance energy transfer (BRET) signal. Addition of small-molecule PKN3 inhibitors that compete with the tracer of the fusion protein in the ATP-binding site results in a detectable loss of BRET signal. The reduction of this BRET signal is the measurement parameter of the NanoBRET assay and determined at an 8-point dose response to determine an in cell target engagement the cellular binding potency.^[37-38]^

**Table 6.**
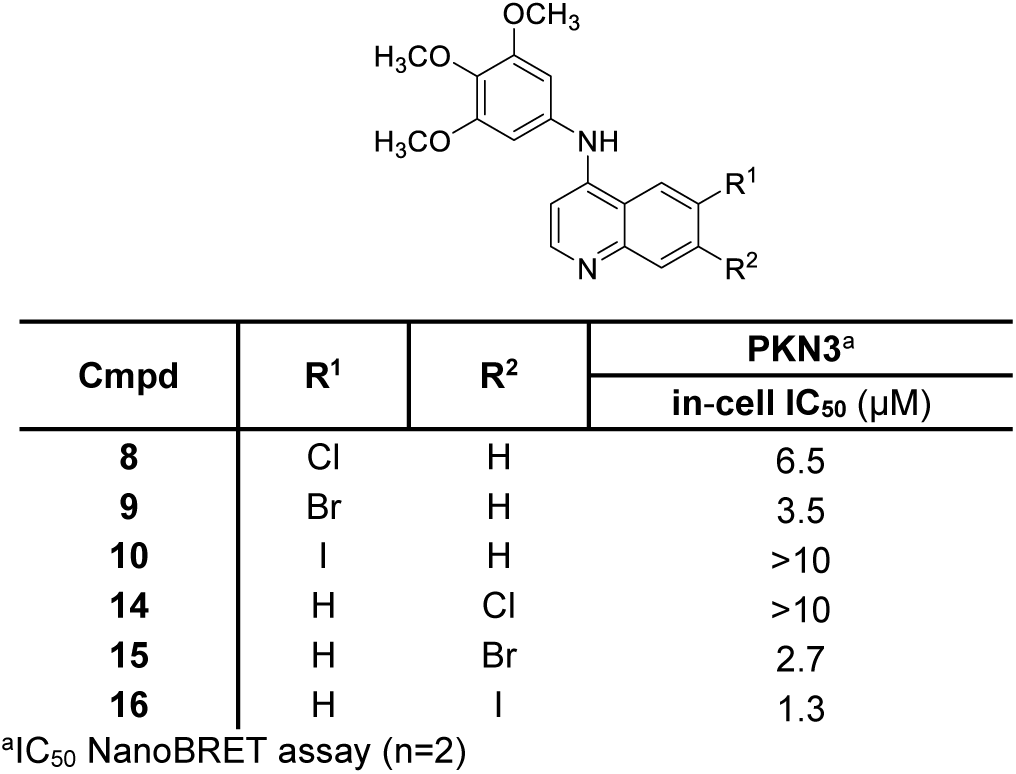
PKN3 NanoBRET results of the most potent analogs.

| Cmpd | R<sup>1</sup> | R<sup>2</sup> | PKN3<sup>a</sup> |
| --- | --- | --- | --- |
|  |  |  | in-cell IC<sub>50</sub> (μM) |
| **8** | Cl | H | 6.5 |
| **9** | Br | H | 3.5 |
| **10** | I | H | >10 |
| **14** | H | Cl | >10 |
| **15** | H | Br | 2.7 |
| **16** | H | I | 1.3 |

<sup>a</sup>IC<sub>50</sub> NanoBRET assay (n=2)

Interestingly, the SLCA and nanoBRET results of this small set of compounds did not have a linear relationship which can partly be explained by cell penetrance among other factors. The most potent compound **9** in the SLCA assay (IC_50_ = 9.3 nM) was not the most potent one in the nanoBRET assay (IC_50_ = 3.5 μM).

The 7-iodo quinoline (**16**) was the most potent derivative with an IC_50_ of 1.3 μM. However, compound **16** revealed significant GAK activity in cells with a bias of just over 17-fold.

In order to model PKN3 we assessed the sequence similarity of studied PKN3 kinase to closely related PKN1 and PKN2. The same kinase family has a high homology (PKN3 vs PKN1 66% and PKN3 vs PKN2 57%), and these kinases are also closely related in the kinome phylogenetic tree.^[39]^ When comparing the experimental x-ray structures of the PKN1 and PKN2 kinase domains, their overall folding is almost identical (Figure S1). This suggests that homology modelling should result in reasonable structural quality. This homology model is especially valuable in since a protein crystal structure of PKN3 is not available. Experimental structures of PKN1 provide information about the apo structure, but also about the binding mode of several co-crystallized inhibitors including lestaurtinib and *bis*indolylmaleimide.^[40]^ The x-ray structure of PRK1 provides information that the HOLO form of both PKN1 and PRK1 that have similar to inhibitor binding modes.^[41]^

Our docking studies using the PKN3 homology model suggest that the quinoline scaffold adopts a similar binding conformation as in EGFR and NAK family kinases as previously reported (Figure 3).^[30]^ The quinoline scaffold binds to the hinge region via H-bond interactions between the amide of Val642 and two weaker aromatic hydrogen bonds to the carbonyl group of Glu640 and Val642. The trimethoxyaniline moiety of **16** accommodates the hydrophobic pocket and forms a a hydrogen bond interaction with the catalytic lysine (Lys588) (Figure 3C).

**Figure 3.**
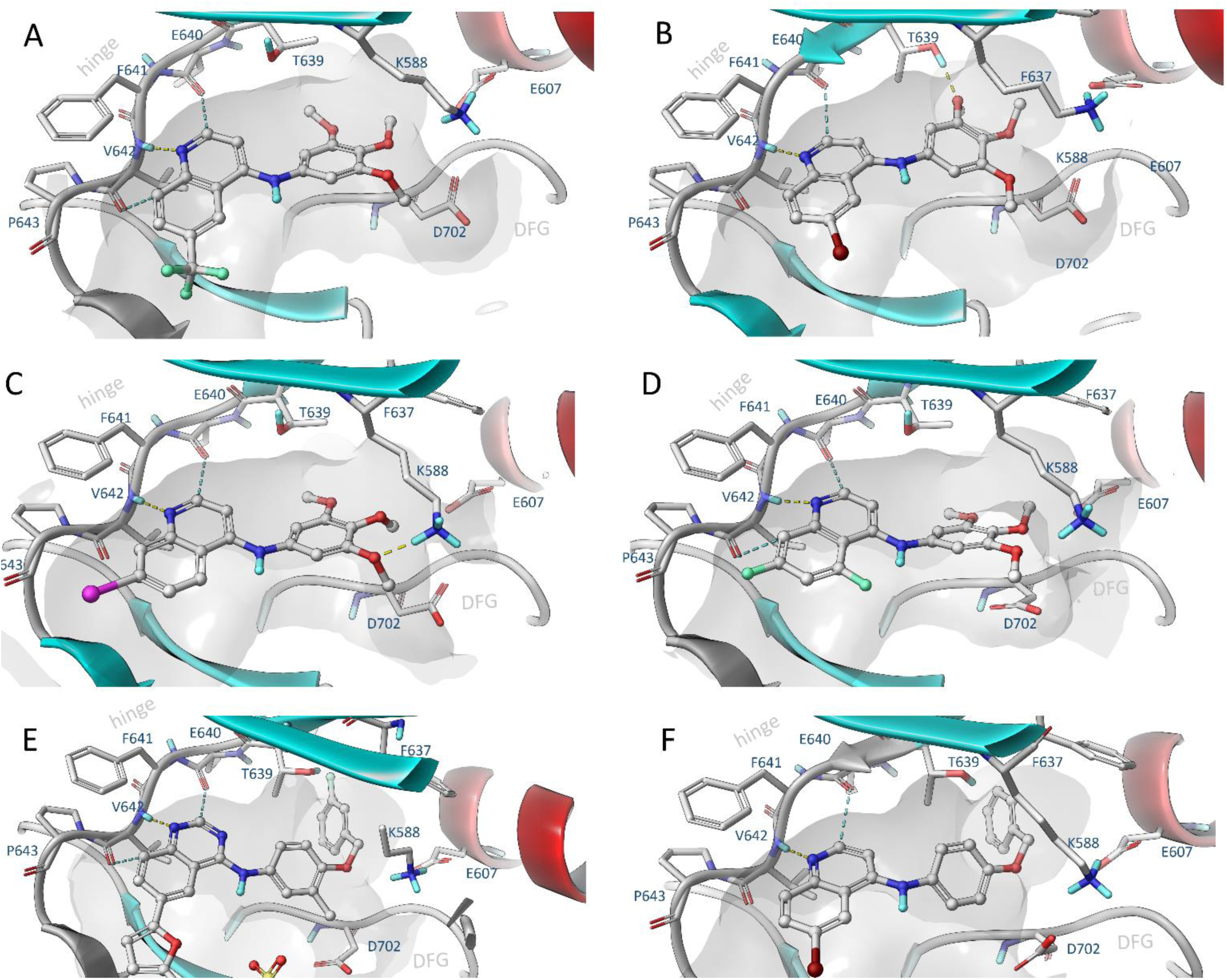
Induced fit docking of selected compounds **2** (A), **9** (B), **16** (C), **39** (D), lapatinib (E) and **50** (F) to PKN3 made by means of comparative modelling using high homologue templates (65%, PDB: 4OTG, 4OTH).

Small-molecule x-ray structures were determined for **52, 54** and **57** (Figure 4).^[42-44]^ This highlighted the influence of the fluorine on the distal part of the aniline ring system.

**Figure 4.**
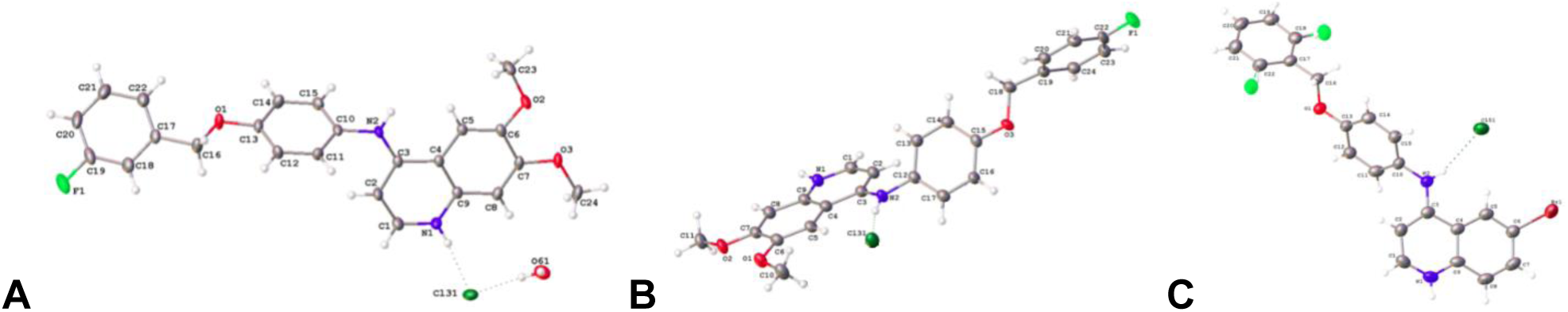
Crystal structures of **52** (A), **54** (B) and **57** (C) with ADP ellipsoids are displayed at 50% probability. Counter ions are shown but solvent molecules are not shown for clarity. Structure A contains two independent molecules per asymmetric unit, only one is shown for clarity.s

## Discussion

Protein kinases present promising drug targets shown by more than 45 inhibitors targeting the ATP binding site of kinases that have been approved for use in the clinic.^[45]^ However, the high clinical efficacy of most of these drugs is based on binding to the conserved ATP binding pocket of more than one kinase.^[46]^ The use of these multi-kinase inhibitors is limited to oncology indications. Inhibitors with significantly improved potency and selectivity profiles are needed for the development of compounds for new treatments outside of oncology.^[47]^ In order to develop those compounds novel approaches are urgently required.

One possible tool amongst others to achieve kinase inhibitors with improved selectivity is the usage of binding assays as a rapid, accurate and robust method to assess potency and potentially wider selectivity of ATP-competitive kinase inhibitors.^[47-48]^ Furthermore, these ligand binding displacement assays provide an accepted direct measurement of kinase inhibition in drug optimization of ATP binding site inhibitors.^[49-50]^ This is a particularly acute point in the case of more neglected kinases such as PKN3 and GAK where there are currently no robust and validated enzyme activity assays available.

In order to not only measure the binding of the synthesized compounds in the isolated enzyme but also in cell the NanoBRET assay presents an excellent technique to measure the target engagement in human cells. Other reported mass spectrometry (MS)-based chemoproteomics approaches rely on the disruption of the plasma membrane and therefore suffer from dissolution of key cellular cofactors like ATP. In contrast, the NanoBRET assay is performed in intact cells. As a result, additionally to be a competitive binder of the ATP binding site the compound needs to be cell permeable in order to be detected by the NanoBRET assay. The cellular thermal shift assay (CETSA) uses intact cells as well. However, CETSA does not quantify the equilibrium-based inhibitor occupancy in contrast to the NanoBRET assay.^[37-38]^ Therefore, the NanoBRET assay was used to determine the in-cell potency of the synthesized PKN3 inhibitors.

Encouraging results from initial PKIS2 screening and literature inhibitors demonstrated the potential of the 4-anilinoquin(az)oline scaffold. We explored several series of substitution patterns around the quin(az)oline core including trimethoxy and hybrid lapatinib derivatives. While strategies were employed to remove key collateral kinase targets including GAK and EGFR they were ultimately only partly successful. The trimethoxy compounds were the most potent compounds on PKN3 with **8, 9, 15** and **16** being the most potent overall in SLCA assay and in cells. We were able to identify **16** as a potent compound against PKN3 with and IC_50_ of 1.3 μM in cells. While several of these quinoline based analogs have been shown to be narrow spectrum across the wider kinome including **2** and **9**. Compound **16** presents a useful potential tool compound to study PKN3 biology. However, further optimization could yield a more potent analog within this series or from the kinase inhibitor landscape.

Our modelling studies with lapatinib and its structure related PKN3 inhibitors suggest that PKN3 may have a more flexible back pocket area. This may give additional structural freedom for further modification of active scaffolds to improve selectivity over GAK and EGFR. However, homology-based docking poses are heavily reflected by structural features derived from template kinases and further structural biology studies with structurally related inhibitors are needed to confirm and validate current docking models.

Taken together this library of compounds provides further evidence that PKN3 can be targeted consistently by a series of small molecule inhibitors. Quinoline-based ATP-competitive kinase inhibitors were designed with high potency *in vitro* and moderate potency in cells. The 4-anilinoquinoline **16** was identified as a potent PKN3 inhibitor with low micromolar potency in cells. The 4-anilinoquinolines series exemplified by **2** and **16** has the potential to yield high-quality chemical probes for use in the elucidation of PKN3 function in cells, and perhaps *in vivo*.

## Experimental Section

### Modelling methods

PKN3 is structurally closely related with other proteins of the serine/threonine protein kinase folding family.^[51]^ The sequence homology to known template structures is feasible 66% - so as a consequence a homology modelling approach was conducted. The PKN3 model was constructed using structure prediction wizard of Maestro 2019.4. At the first stage a blast search was carried out in order to find suitable template structures. Sequences taken from crystal structures of PRK1 Catalytic Domain (PDB: 4OTD, 4OTG, 4OTH, 4OTI) and Human Protein Kinase N2 (PDB: 4CRS) were used for pairwise sequence alignment using clustalW methodology, which is suitable in this case, due to the structural similarity. High resolution structure, PDB:4OTH was selected as template for comparative modelling (identity 66%, positives 80%, alignment length 337 (kinase domain), resolution 1.8Å). Due to the good sequence similarity, a homology model was constructed using knowledge-based approach of the wizard employing multi template approach, where the model was mainly constructed based on PDB:4OTH, including ligand occupying the ATP binding pocket. The missing activation loop of PDB:4OTH was modeled based on other PDB structure of the same template (PDB:4OTG). Loop refinement was run to optimize the structure of *C*-terminal end.

Prior to docking studies, the homology model of the PKN3 structure was pre-processed by using the protein preparation wizard tool of Schrödinger Suite 2019-4 (Protein Preparation Wizard uses modules: Epik; Impact and Prime, Schrödinger, LLC, New York, NY, 2019). Structures of small molecule ligands were parametrized and minimized using Ligprep module (LigPrep, Schrödinger, LLC, New York, NY, 2019.). Molecular docking studies were computed using Induced fit workflow of Schrödinger employing SP-setting for Glide docking module and side chain 5Å were consider for conformational refinement using Prime module. Hydrogen bond constraint was added to hinge residue VAL642 to improved convergence of docking poses.Results were consistent with co-crystal binding modes in other kinases, eg GAK and the 4-anilinoquin(az)oline gefitinib.^[52]^

### Biology

#### SLCA Method

The split-luciferase kinase assays were performed by Luceome Biotechnologies. Briefly, split-firefly luciferase constructs were translated and assayed in cell-free system according to literature protocol.^[29, 30]^ Data were collected via a homogeneous competition binding assay where the displacement of an active site dependent probe by an inhibitor is measured by a change in luminescence signal. Testing was performed with either DMSO (no-inhibitor control) or a compound solution in DMSO, followed by incubation in the presence of a kinase specific probe. Luminescence was measured upon addition of a luciferin assay reagent to each assay solution. All PKIS2 library members and SAR analogs were screened at 1 µM in duplicate against PKN3. IC_50_ values (8-point dose response format) were determined by plotting percent activity remaining against inhibitor concentration.

#### NanoBRET method. Cell Transfections and BRET Measurements

*N*-terminal NanoLuc/Kinase fusions were encoded in pFN31K or pFC32K expression vectors (Promega), including flexible Gly-Ser-Ser-Gly linkers between Nluc and each full-length kinase. For cellular BRET target engagement experiments, HEK-293 cells were transfected with NLuc/target fusion constructs using FuGENE HD (Promega) according to the manufacturer’s protocol. Briefly, Nluc/target fusion constructs were diluted into Opti-MEM followed by Transfection Carrier DNA (Promega) at a mass ratio of 1:10 (mass/mass), after which FuGENE HD was added at a ratio of 1:3 (mg DNA: mL FuGENE HD). 1 part (vol) of FuGENE HD complexes thus formed were combined with 20 parts (vol) of HEK-293 cells in DMEM with 10% FBS plated at a density of 2 × 10^5^ per mL into 96-well plates (Corning), followed by incubation in a humidified, 37°C/5% CO_2_ incubator for 20-30 hr. BRET assays were performed in white, 96-well cell culture treated plates (Corning) at a density of 2 × 104 cells/well.

Following transfection, DMEM was exchanged for Opti-MEM. All chemical inhibitors were prepared as concentrated stock solutions in DMSO (Sigma-Aldrich) and diluted in Opti-MEM (unless otherwise noted) to prepare working stocks. Cells were equilibrated for 2 h with energy transfer probes and test compound prior to BRET measurements. Energy transfer probes were prepared at a working concentration of 20X in tracer dilution buffer (12.5 mM HEPES, 31.25% PEG-400, pH 7.5). For target engagement analysis, the energy transfer probes were added to the cells at concentrations optimized for each target. For analysis of PKN3-NL, energy transfer probe K-5 was used at a final concentration of 1000 nM. To measure BRET, NanoBRET NanoGlo Substrate and Extracellular NanoLuc Inhibitor (Promega) were added according to the manufacturer’s recommended protocol, and filtered luminescence was measured on a GloMax Discover luminometer equipped with 450 nM BP filter (donor) and 600 nM LP filter (acceptor), using 0.5 s integration time. Milli-BRET units (mBU) are calculated by multiplying the raw BRET values by 1000. Competitive displacement data were plotted with GraphPad Prism software and data were fit to Equation 1 [log(inhibitor) vs. response -- Variable slope (four parameters)] to determine the IC_50_ value; Y = Bottom + (Top-Bottom)/(1 + 10^((LogIC_50_-X)*HillSlope)). (Equation 1).

If the curve was insufficiently described by the raw data, then they were normalized to the controls-BRET in the presence of only energy transfer probe, and BRET in the absence of the energy transfer probe and test compound-then plotted to fit Equation 2 [log(inhibitor) vs. normalized response -- Variable slope] to determine the IC_50_ value; Y=100/(1+10^((LogIC_50_-X)*HillSlope))). (Equation 2). For all BRET data shown, no individual data points were omitted.

### Chemistry

#### General Procedure

4-chloroquinoline derivative (150 mg, 0.67 mmol) and aniline (0.74 mmol) was suspended in ethanol (10 mL) and heated to reflux for 16 h. The crude mixture was purified by flash chromatography using EtOAc/hexane followed by 1–5% methanol in EtOAc (or by re-crystallization). After solvent removal under reduced pressure, the product was obtained. All compounds were *>*98% pure by ^1^H/^13^C NMR and LCMS. Compounds **2** and **4**-**39** were prepared as previously described.^[35,36, 53]^

Automated flash chromatography: Isolera^™^ Prime, Version 3.0 (Biotage^®^); RediSep^®^ Rf columns (Teledyne ISCO^™^), normal-phase, prepacked, 5g; parentheses include: cartridge size, flow rate, eluent, fractions size was always 12 mL. The crude mixture was purified by automatic flash chromatography (5g column, flowrate: 6 ml/min, *n*-hexane → *n*-hexane : ethyl acetate 80:20 → ethyl acetate → ethyl acetate : methanol = 90:10) and the solvent was evaporated under reduced pressure to yield the title compound.

#### *N*-(4-Ethoxyphenyl)-6,7-dimethoxyquinolin-4-amine (40)

prepared as decribed with an additional washing step of ether (10 mL) to cause the precipitation of the crude product prior to filtration to afford a mustard solid (198 mg, 0.610 mmol, 91 %); m.p. 241 °C; ^1^H NMR (400 MHz, [D_6_]DMSO) δ 10.60 (s, 1H), 8.28 (d, *J* = 6.9 Hz, 1H), 8.16 (s, 1H), 7.46 (s, 1H), 7.40 – 7.31 (m, 2H), 7.13 – 7.04 (m, 2H), 6.54 (d, *J* = 6.9 Hz, 1H), 4.08 (q, *J* = 6.9 Hz, 2H), 3.99 (s, 3H), 3.96 (s, 3H), 1.36 ppm (t, *J* = 6.9 Hz, 3H); ^13^C NMR (101 MHz, [D_6_]DMSO) δ 157.4, 154.4, 153.5, 149.3, 139.8, 135.4, 129.9, 127.2, 115.5, 111.3, 102.7, 100.0, 98.7, 63.4, 56.7, 56.1, 14.6 ppm; HRMS-ESI (m/z): [M+H]^+^ calcd for C_19_H_21_N_2_O_3_ ^+^ 325.1547, found 325.1543; LC: t_R_ = 3.84 min, purity > 98 %.

#### 6-Bromo-*N*-(4-ethoxyphenyl)quinolin-4-amine (41)

was obtained as a light brown solid (197 mg, 0.572 mmol, 93 %); m.p. 151 °C; ^1^H NMR (400 MHz, [D_6_]DMSO) δ 10.98 (s, 1H), 9.13 (d, *J* = 2.0 Hz, 1H), 8.48 (d, *J* = 7.0 Hz, 1H), 8.15 (dd, *J* = 9.0, 2.0 Hz, 1H), 8.05 (d, *J* = 9.0 Hz, 1H), 7.40 – 7.34 (m, 2H), 7.14 – 7.08 (m, 2H), 6.68 (d, *J* = 7.0 Hz, 1H), 4.09 (q, *J* = 7.0 Hz, 2H), 1.36 ppm (t, *J* = 7.0 Hz, 3H). ^13^C NMR (101 MHz, [D_6_]DMSO) δ 157.7, 154.4, 142.8, 137.3, 136.5, 129.3, 127.0, 126.1, 122.4, 119.7, 118.4, 115.6, 100.1, 63.4, 14.6 ppm; HRMS-ESI (m/z): [M+H]^+^ calcd for C_17_H_16_BrN_2_O^+^ 343.0441, found 343.0440; LC: t_R_ = 3.97 min, purity > 98 %.

#### *N*-(4-Ethoxyphenyl)-5,7-difluoroquinolin-4-amine (42)

was prepared on a larger scale (4-chloro-5,7-difluoroquinoline (500 mg, 2.51 mmol) and aniline (2.76 mmol) and additional following washing steps with ethyl acetate and a mixture of ethyl acetate/methanol 9:1) to afford a bright yellow solid (153 mg, 0.509 mmol, 20 %); m.p. 238 °C; ^1^H NMR (400 MHz, [D_6_]DMSO) δ 10.22 (d, *J* = 11.8 Hz, 1H), 8.42 (d, *J* = 7.2 Hz, 1H), 7.79 (dd, *J* = 12.7, 9.1 Hz, 2H), 7.38 – 7.32 (m, 2H), 7.14 – 7.08 (m, 2H), 6.50 (d, *J* = 7.1 Hz, 1H), 4.09 (q, *J* = 6.9 Hz, 2H), 1.36 ppm (t, *J* = 6.9 Hz, 3H); ^13^C NMR (101 MHz, [D_6_]DMSO) δ 164.1 (dd, *J* = 253.3, 15.6 Hz), 160.7 (dd, *J* = 254.2, 15.3 Hz), 158.6, 155.3 (d, *J* = 3.3 Hz), 143.5, 141.4 (dd, *J* = 15.1, 6.5 Hz), 129.9, 128.4, 116.1, 105.67 (dd, *J* = 13.4, 2.2 Hz), 104.3 – 103.2 (m), 102.1 (dd, *J* = 25.3, 4.5 Hz), 101.4, 63.9, 15.1 ppm. HRMS-ESI (m/z): [M+H]^+^ calcd for C_17_H_15_F_2_N_2_O^+^ 301.1147, found 301.1147; LC: t_R_ = 3.69 min, purity > 98 %.

#### *N*-(4-Isopropoxyphenyl)-6,7-dimethoxynaphthalen-1-amine (43)

was obtained as a light yellow solid (195 mg, 0.578 mmol, 86 %); m.p. 209 °C; ^1^H NMR (400 MHz, [D_6_]DMSO) δ 10.52 (s, 1H), 8.28 (d, *J* = 6.9 Hz, 1H), 8.13 (s, 1H), 7.44 (s, 1H), 7.39 – 7.30 (m, 2H), 7.11 – 7.03 (m, 2H), 6.56 (d, *J* = 6.9 Hz, 1H), 4.67 (hept, *J* = 6.0 Hz, 1H), 3.99 (s, 3H), 3.96 (s, 3H), 1.31 (s, 3H), 1.30 ppm (s, 3H); ^13^C NMR (101 MHz, [D_6_]DMSO) δ 156.3, 154.3, 153.4, 149.2, 140.1, 135.7, 129.8, 127.2, 116.6, 111.4, 102.6, 100.3, 98.8, 69.5, 56.6, 56.1, 21.8 ppm; HRMS-ESI (m/z): [M+H]^+^ calcd for C_21_H_24_NO_3_ ^+^ 338.4265, found 339.1700; LC: t_R_ = 4.04 min, purity > 98 %.

#### 6-Bromo-*N*-(4-isopropoxyphenyl)quinolin-4-amine (44)

was obtained as a light brown solid (107 mg, 0.299 mmol, 48 %); m.p. 149 °C; ^1^H NMR (400 MHz, [D_6_]DMSO) δ 10.86 (s, 1H), 9.09 (d, *J* = 2.0 Hz, 1H), 8.45 (d, *J* = 6.9 Hz, 1H), 8.12 (dd, *J* = 9.0, 2.0 Hz, 1H), 8.02 (d, *J* = 9.0 Hz, 1H), 7.37 – 7.31 (m, 2H), 7.10 – 7.04 (m, 2H), 6.68 (d, *J* = 6.9 Hz, 1H), 4.66 (hept, *J* = 6.0 Hz, 1H), 1.30 (s, 3H), 1.28 ppm (s, 3H); ^13^C NMR (101 MHz, [D_6_]DMSO) δ 156.6, 154.1, 143.2, 137.8, 136.3, 129.3, 126.9, 126.0, 122.9, 119.6, 118.5, 116.6, 100.2, 69.6, 21.8 ppm; HRMS-ESI (m/z): [M+H]^+^ calcd for C_18_H_18_BrN_2_O^+^ 357.0597, found 357.0603; LC: t_R_ = 4.55 min, purity > 98 %.

#### 6-Bromo-*N*-(4-cyclobutoxyphenyl)quinolin-4-amine (45)

was prepared on a larger scale (6-bromo-4-chloroquinoline (500 mg, 2.51 mmol) and aniline (2.76 mmol)) and additional following washing steps with ethyl acetate and a mixture of ethyl acetate/methanol 9:1) to afford a light yellow solid (232 mg, 0.738 mmol, 29 %); m.p. 236 °C; ^1^H NMR (400 MHz, [D_6_]DMSO) δ 10.21 (d, *J* = 11.8 Hz, 1H), 8.42 (d, *J* = 7.2 Hz, 1H), 7.85 – 7.74 (m, 2H), 7.37 – 7.31 (m, 2H), 7.13 – 7.07 (m, 2H), 6.52 (d, *J* = 7.1 Hz, 1H), 4.68 (hept, *J* = 6.0 Hz, 1H), 1.31 (s, 3H), 1.30 ppm (s, 3H); ^13^C NMR (101 MHz, [D_6_]DMSO) δ 163.6 (dd, *J* = 253.3, 15.4 Hz), 160.3 (dd, *J* = 257.9, 15.3 Hz), 157.1, 154.8 (d, *J* = 3.3 Hz), 143.1, 140.9 (dd, *J* = 15.0, 6.6 Hz), 129.3, 127.9, 116.7, 105.2 (dd, *J* = 13.1, 2.2 Hz), 103.2 (dd, *J* = 28.9, 26.8 Hz), 101.6 (dd, *J* = 25.1, 4.3 Hz), 101.0, 69.6, 21.8 ppm; HRMS-ESI (m/z): [M+H]^+^ calcd for C_18_H_17_F_2_N_2_O^+^ 315.1303, found 315.1300; LC: t_R_ = 3.90 min, purity > 98 %.

#### *N*-(4-Cyclobutoxyphenyl)-6,7-dimethoxyquinolin-4-amine (46)

was obtained as a light yellow solid (170 mg, 0.485 mmol, 72 %); m.p. 234 °C; ^1^H NMR (400 MHz, [D_6_]DMSO) δ 10.50 (s, 1H), 8.27 (d, *J* = 6.8 Hz, 1H), 8.13 (s, 1H), 7.44 (s, 1H), 7.38 – 7.30 (m, 2H), 7.04 – 6.96 (m, 2H), 6.54 (d, *J* = 6.8 Hz, 1H), 4.73 (p, *J* = 7.0 Hz, 1H), 3.98 (s, 3H), 3.95 (s, 3H), 2.49 – 2.41 (m, 1H), 2.14 – 2.01 (m, 2H), 1.87 – 1.75 (m, 1H), 1.73 – 1.59 ppm (m, 1H); ^13^C NMR (101 MHz, [D_6_]DMSO) δ 155.9, 154.4, 153.5, 149.3, 139.9, 135.5, 130.0, 127.2, 115.8, 111.3, 102.7, 100.1, 98.8, 71.0, 56.7, 56.1, 30.1, 12.8 ppm; HRMS-ESI (m/z): [M+H]^+^ calcd for C_21_H_23_N_2_O_3_ ^+^ 351.1703, found 351.1697; LC: t_R_ = 4.22 min, purity > 98 %.

#### 6-Bromo-*N*-(4-cyclobutoxyphenyl)quinolin-4-amine (47)

was obtained as a mustard solid (144 mg, 0.390 mmol, 46 %); m.p. > 250 °C; ^1^H NMR (400 MHz, [D_6_]DMSO) δ 10.98 (s, 1H), 9.14 (d, *J* = 1.8 Hz, 1H), 8.47 (d, *J* = 6.9 Hz, 1H), 8.15 (dd, *J* = 9.0, 1.8 Hz, 1H), 8.06 (d, *J* = 9.0 Hz, 1H), 7.40 – 7.33 (m, 2H), 7.06 – 6.98 (m, 2H), 6.68 (d, *J* = 6.9 Hz, 1H), 4.74 (p, *J* = 7.1 Hz, 1H), 2.49 – 2.41 (m, 2H), 2.14 – 2.00 (m, 2H), 1.87 – 1.75 (m, 1H), 1.75 – 1.58 ppm (m, 1H); ^13^C NMR (101 MHz, [D_6_]DMSO) δ 156.3, 154.3, 142.9, 137.4, 136.4, 129.5, 127.0, 126.1, 122.5, 119.6, 118.4, 115.9, 100.1, 71.0, 30.1, 12.8 ppm; calcd for C_19_H_18_BrN_2_O^+^ 369.0597, found 369.0594; LC: t_R_ = 4.33 min, purity > 98 %.

#### 6-Bromo-*N*-(4-cyclobutoxyphenyl)quinolin-4-amine (48)

was obtained as a tan solid (58 mg, 0.178 mmol, 24 %); m.p. 238 °C; ^1^H NMR (400 MHz, [D_6_]DMSO) δ 10.12 (d, *J* = 10.4 Hz, 1H), 8.41 (d, *J* = 7.1 Hz, 1H), 7.82 – 7.71 (m, 2H), 7.37 – 7.29 (m, 2H), 7.05 – 6.99 (m, 2H), 6.50 (d, *J* = 7.1 Hz, 1H), 4.75 (p, *J* = 7.1 Hz, 1H), 2.49 – 2.41 (m, 2H), 2.13 – 2.00 (m, 3H), 1.87 – 1.75 (m, 1H), 1.73 – 1.59 ppm (m, 1H); ^13^C NMR (101 MHz, [D_6_]DMSO) δ 163.5 (dd, *J* = 253.0, 15.5 Hz), 160.2 (dd, *J* = 257.8, 15.4 Hz), 156.6, 154.6 (d, *J* = 3.4 Hz), 143.6, 141.4 (dd, *J* = 15.0, 6.2 Hz), 129.7, 127.9, 116.0, 105.3 (dd, *J* = 13.0, 2.0 Hz), 103.8 – 102.7 (m), 102.0 (dd, *J* = 24.3, 3.3 Hz), 101.0, 71.0, 30.0, 12.8 ppm; HRMS-ESI (m/z): [M+H]^+^ calcd for C_19_H_17_F_2_N_2_O^+^ 327.1303, found 327.1299; LC: t_R_ = 4.09 min, purity > 98 %.

#### *N*-(4-(benzyloxy)phenyl)-6,7-dimethoxyquinolin-4-amine (49)

was obtained as a beige solid (187 mg, 0.483 mmol, 72%). MP >250 °C; ^1^H NMR (400 MHz, [D_6_]DMSO) δ 10.68 (s, 1H), 8.28 (d, *J* = 7.0 Hz, 1H), 8.18 (s, 1H), 7.78 – 7.23 (m, 8H), 7.23 – 7.12 (m, 2H), 6.55 (d, *J* = 7.0 Hz, 1H), 5.17 (s, 2H), 3.99 (s, 3H), 3.95 (s, 3H). ^13^C NMR (101 MHz, [D_6_]DMSO) δ 157.2, 154.4, 153.6, 149.3, 139.6, 136.8, 135.2, 130.2, 128.5 (2C, s), 127.9, 127.8 (2C, s), 127.2, 115.9, 111.3, 102.8, 99.8, 98.7, 69.6, 56.7, 56.1. calcd for C_24_H_23_N_2_O_3_ ^+^ 387.1703, found 387.1707; LC: t_R_ = 4.89 min, purity > 98 %.

#### *N*-(4-(benzyloxy)phenyl)-6-bromoquinolin-4-amine (50)

Was obtained as a yellow solid (183 mg, 0.452 mmol, 73%). MP 129-131 °C; ^1^H NMR (400 MHz, [D_6_]DMSO) δ 10.96 (s, 1H), 9.11 (d, *J* = 2.1 Hz, 1H), 8.48 (d, *J* = 7.0 Hz, 1H), 8.16 (dd, *J* = 9.0, 2.0 Hz, 1H), 8.04 (d, *J* = 9.0 Hz, 1H), 7.72 – 7.25 (m, 7H), 7.25 – 7.16 (m, 2H), 6.69 (d, *J* = 7.0 Hz, 1H), 5.18 (s, 2H). ^13^C NMR (101 MHz, [D_6_]DMSO) δ 157.5, 154.4, 142.9, 137.3, 136.8, 136.5, 129.6, 128.5 (2C, s), 128.0, 127.8 (2C, s), 127.0, 126.1, 122.5, 119.7, 118.4, 116.0, 100.1, 69.6. calcd for C_22_H_18_BrN_2_O^+^ 405.0597, found 405.0603; LC: t_R_ = 5.03 min, purity > 98 %.

#### *N*-(4-(Benzyloxy)phenyl)-5,7-difluoroquinolin-4-amine (51)

was obtained as a mustard solid (130 mg, 0.359 mmol, 58 %); m.p. 231 °C; ^1^H NMR (400 MHz, [D_6_]DMSO) δ 10.22 (d, *J* = 12.0 Hz, 1H), 8.43 (d, *J* = 7.1 Hz, 1H), 7.84 – 7.73 (m, 2H), 7.52 – 7.46 (m, 2H), 7.45 – 7.32 (m, 5H), 7.25 – 7.18 (m, 2H), 6.50 (d, *J* = 7.1 Hz, 1H), 5.18 ppm (s, 2H); ^13^C NMR (101 MHz, [D_6_]DMSO) δ 163.63 (dd, *J* = 253.6, 15.3 Hz), 160.25 (dd, *J* = 258.0, 15.2 Hz), 157.88, 154.82 (d, *J* = 2.9 Hz), 143.11, 140.91 (dd, *J* = 15.0, 6.4 Hz), 136.75, 129.82, 128.49, 127.96, 127.93, 127.77, 116.05, 105.22 (dd, *J* = 13.1, 2.1 Hz), 103.74 – 102.64 (m), 101.66 (dd, *J* = 25.1, 4.2 Hz), 100.99, 69.53 ppm; HRMS-ESI (m/z): [M+H]^+^ calcd for C_22_H_17_F_2_N_2_O^+^ 363.1303, found 363.1298; LC: t_R_ = 4.32 min, purity > 98 %.

#### *N*-(4-((3-fluorobenzyl)oxy)phenyl)-6,7-dimethoxyquinolin-4-amine (52)

was obtained as a dark green solid (130 mg, 0.321 mmol, 48%). MP decomp. 220 °C; ^1^H NMR (400 MHz, [D_6_]DMSO) δ 10.65 (s, 1H), 8.29 (d, *J* = 6.9 Hz, 1H), 8.17 (s, 1H), 7.50 – 7.43 (m, 2H), 7.42 – 7.36 (m, 2H), 7.35 – 7.28 (m, 2H), 7.22 – 7.10 (m, 3H), 6.55 (d, *J* = 6.9 Hz, 1H), 5.20 (s, 2H), 3.99 (s, 3H), 3.95 (s, 3H). ^13^C NMR (101 MHz, [D_6_]DMSO) δ 162.2 (d, *J* = 243.7 Hz), 156.9, 154.4, 153.5, 149.3, 139.8, 139.8 (d, *J* = 1.6 Hz), 135.3, 130.5 (d, *J* = 8.3 Hz), 130.4, 127.2, 123.6 (d, *J* = 2.7 Hz), 115.9, 114.7 (d, *J* = 20.9 Hz), 114.3 (d, *J* = 21.8 Hz), 111.4, 102.7, 100.0, 98.8, 68.7 (d, *J* = 2.1 Hz), 56.7, 56.1 ppm; HRMS-ESI (m/z): [M+H]^+^ calcd for C_24_H_22_FN_2_O_3_ ^+^ 405.1609, found 405.0617, ; LC: t_R_ = 4.46 min, purity > 98 %.

#### 6-bromo-*N*-(4-((3-fluorobenzyl)oxy)phenyl)quinolin-4-amine(53)

was obtained as a dark green solid (144 mg, 0.340 mmol, 55%). MP 111-113 °C; ^1^H NMR (400 MHz, [D_6_]DMSO) δ 11.01 (s, 1H), 9.13 (d, *J* = 2.0 Hz, 1H), 8.48 (d, *J* = 7.0 Hz, 1H), 8.15 (dd, *J* = 9.0, 2.0 Hz, 1H), 8.06 (d, *J* = 9.0 Hz, 1H), 7.47 (td, *J* = 8.1, 6.0 Hz, 1H), 7.43 – 7.37 (m, 2H), 7.36 – 7.28 (m, 2H), 7.24 – 7.19 (m, 2H), 7.17 (ddd, *J* = 8.2, 3.1, 1.1 Hz, 1H), 6.69 (d, *J* = 7.0 Hz, 1H), 5.21 (s, 2H). ^13^C NMR (101 MHz, [D_6_]DMSO) δ 162.2 (d, *J* = 243.5 Hz), 157.3, 154.4, 142.8, 139.7 (d, *J* = 7.3 Hz), 137.3, 136.5, 130.5 (d, *J* = 8.3 Hz), 129.8, 127.0 (2C, s), 126.1, 123.6 (d, *J* = 2.7 Hz), 122.4, 119.7, 118.4, 116.0 (2C, s), 114.7 (d, *J* = 20.9 Hz), 114.3 (d, *J* = 21.9 Hz), 100.1, 68.7 ppm (d, *J* = 1.9 Hz); HRMS-ESI (m/z): [M+H]^+^ calcd for C_22_H_17_BrFN_2_O^+^ 423.0503, found 423.0512; LC: t_R_ = 5.11 min, purity > 98 %.

#### *N*-(4-((4-fluorobenzyl)oxy)phenyl)-6,7-dimethoxyquinolin-4-amine (54)

was obtained as a grey solid (176 mg, 0.426 mmol, 65%). MP decomp. 210 °C; ^1^H NMR (400 MHz, [D_6_]DMSO) δ 10.71 (s, 1H), 8.27 (d, *J* = 7.0 Hz, 1H), 8.19 (s, 1H), 7.57 – 7.49 (m, 2H), 7.47 (s, 1H), 7.42 – 7.35 (m, 2H), 7.28 – 7.20 (m, 2H), 7.20 – 7.13 (m, 2H), 6.54 (d, *J* = 6.9 Hz, 1H), 5.14 (s, 2H), 3.99 (s, 3H), 3.94 (s, 3H). ^13^C NMR (101 MHz, [D_6_]DMSO) δ 161.8 (d, *J* = 243.8 Hz), 157.0, 154.4, 153.5, 149.2, 139.6, 135.3, 133.1 (d, *J* = 3.0 Hz), 130.3, 130.0 (d, *J* = 8.3 Hz, 2C), 127.2 (s, 2C), 115.9 (s, 2C), 115.3 (d, *J* = 21.4 Hz, 2C), 111.4, 102.8, 99.9, 98.7, 68.8, 56.8, 56.1. HRMS-ESI (m/z): [M+H]^+^ calcd for C_24_H_22_FN_2_O_3_ ^+^ 405.1609, found 405.1613; LC: t_R_ = 4.91 min, purity > 98 %.

#### 6-bromo-*N*-{4-[(4-fluorophenyl)methoxy]phenyl}quinolin-4-amine (55)

was obtained as a dark yellow solid (178 mg, 0.421 mmol, 68%). MP 128-120 °C; ^1^H NMR (400 MHz, [D_6_]DMSO) δ 10.95 (s, 1H), 9.11 (d, *J* = 2.0 Hz, 1H), 8.48 (d, *J* = 7.0 Hz, 1H), 8.15 (dd, *J* = 9.0, 2.0 Hz, 1H), 8.04 (d, *J* = 9.0 Hz, 1H), 7.71 – 7.47 (m, 2H), 7.47 – 7.32 (m, 2H), 7.32 – 7.00 (m, 4H), 6.69 (d, *J* = 6.9 Hz, 1H), 5.16 (s, 2H). ^13^C NMR (101 MHz, [D_6_]DMSO) δ 161.8 (d, *J* = 243.5 Hz), 157.4, 154.3, 143.0, 137.4, 136.4, 133.0 (d, *J* = 3.0 Hz), 130.1 (d, *J* = 8.2 Hz, 2C), 129.7, 127.0 (s, 2C), 126.1, 122.6, 119.7, 118.4, 116.0 (s, 2C), 115.3 (d, *J* = 21.4 Hz, 2C), 100.1, 68.8 ppm; HRMS-ESI (m/z): [M+H]^+^ calcd for C_22_H_17_BrFN_2_O^+^ 423.0503, found 423.0511; LC: t_R_ = 5.09 min, purity > 98 %.

#### *N*-(4-((2-fluorobenzyl)oxy)phenyl)-6,7-dimethoxyquinolin-4-amine (56)

was obtained as a grey solid (94 mg, 0.235 mmol, 35%). MP 232-234 °C; ^1^H NMR (400 MHz, [D_6_]DMSO) δ 10.56 (s, 1H), 8.29 (d, *J* = 6.9 Hz, 1H), 8.14 (s, 1H), 7.60 (td, *J* = 7.6, 1.7 Hz, 1H), 7.56 – 7.42 (m, 2H), 7.42 – 7.32 (m, 2H), 7.32 – 7.23 (m, 2H), 7.23 – 7.07 (m, 2H), 6.57 (d, *J* = 6.9 Hz, 1H), 5.20 (s, 2H), 3.99 (s, 3H), 3.96 (s, 3H). ^13^C NMR (101 MHz, [D_6_]DMSO) δ 160.5 (d, *J* = 246.2 Hz), 157.0, 154.4, 153.4, 149.3, 140.0, 135.6, 130.8 (d, *J* = 4.1 Hz), 130.6, 130.5, 127.2 (2C, s), 124.6 (d, *J* = 3.5 Hz), 123.6 (d, *J* = 14.5 Hz), 115.9 (2C, s), 115.4 (d, *J* = 21.0 Hz), 111.4, 102.6, 100.2, 98.8, 63.9 (d, *J* = 3.6 Hz), 56.7, 56.1 ppm; HRMS-ESI (m/z): [M+H]^+^ calcd for C_24_H_22_FN_2_O_3_ ^+^ 405.1609, found 405.1612; LC: t_R_ = 4.90 min, purity > 98 %.

#### 6-bromo*-N*-(4-((2-fluorobenzyl)oxy)phenyl)quinolin-4-amine(57)

was obtained as a green solid (113 mg, 0.266 mmol, 43%). MP 126-128 °C; ^1^H NMR (400 MHz, [D_6_]DMSO) δ 10.89 (s, 1H), 9.08 (d, *J* = 2.1 Hz, 1H), 8.49 (d, *J* = 7.0 Hz, 1H), 8.16 (dd, *J* = 9.0, 2.0 Hz, 1H), 8.02 (d, *J* = 9.0 Hz, 1H), 7.60 (td, *J* = 7.5, 1.7 Hz, 1H), 7.49 – 7.22 (m, 6H), 6.71 (d, *J* = 7.0 Hz, 1H), 5.21 (s, 2H). ^13^C NMR (101 MHz, [D_6_]DMSO) δ 160.5 (d, *J* = 246.2 Hz), 157.4, 154.3, 143.1, 137.4, 136.5, 130.9 (d, *J* = 4.0 Hz), 130.6 (d, *J* = 8.3 Hz), 129.9, 127.0 (2C, s), 126.0, 124.6 (d, *J* = 3.4 Hz), 123.5 (d, *J* = 14.5 Hz), 122.6, 119.7, 118.4, 116.0 (2C, s), 115.5 (d, *J* = 20.9 Hz), 100.2, 64.0 ppm (d, *J* = 3.6 Hz); HRMS-ESI (m/z): [M+H]^+^ calcd for C_22_H_17_BrFN_2_O^+^ 423.0503, found 423.0512; LC: t_R_ = 5.07 Min, purity > 98 %.

## Acknowledgements

The SGC is a registered charity (number 1097737) that receives funds from AbbVie, Bayer Pharma AG, Boehringer Ingelheim, Canada Foundation for Innovation, Eshelman Institute for Innovation, Genome Canada, Innovative Medicines Initiative (EU/EFPIA) [ULTRA-DD grant no. 115766], Janssen, Merck KGaA Darmstadt Germany, MSD, Novartis Pharma AG, Ontario Ministry of Economic Development and Innovation, Pfizer, São Paulo Research Foundation-FAPESP, Takeda, and Wellcome [106169/ZZ14/Z]. Research reported in this publication was also supported by a grant from the National Center for Advancing Translational Sciences (NCATS) of the National Institute of Health “Tools for Accelerating R&D for Historically Understudied Protein Kinases” (1R44TR001916-01) and by the National Institute of Health “Illuminating the Druggable Genome program” (grant number 1U24DK116204-01). The content is solely the responsibility of the authors and does not necessarily represent the official views of the National Institute of Health. In addition, we thank Biocenter Finland/DDCB for financial support and CSC – IT Center for Science Ltd. Finland for the use of their facilities, software licenses and computational resources. We are grateful Dr. Brandie Ehrmann and Diane Wallace for LC-MS/HRMS support provided by the Mass Spectrometry Core Laboratory at the University of North Carolina at Chapel Hill. We also thank the EPSRC UK National Crystallography Service for funding and collection of the crystallographic data for **52, 54** and **57**.

## Graphic

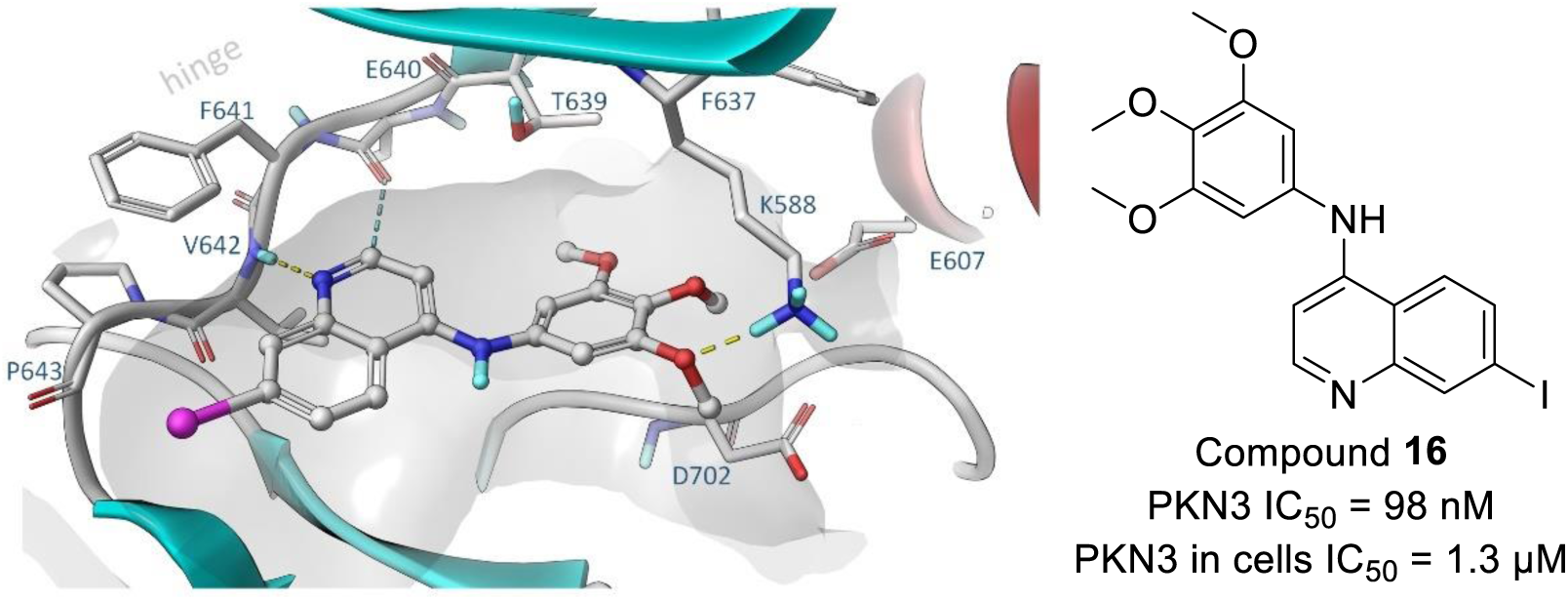

We developed several small focused libraries of 4-anilinoquinolines to investigate a screening hit on a understudied kinase Protein Kinase Novel 3 (PKN3). This to the identification of 7-iodo-*N*-(3,4,5-trimethoxyphenyl)quinolin-4-amine **16** as a potent inhibitor of PKN3 with an IC_50_ of 1.3 μM in cells. Compound **16** is a potentially useful tool to study PKN3 biology including links to pancreatic and prostate cancer, along with T-cell acute lymphoblastic leukemia. These compounds may be useful tools to explore the therapeutic potential of PKN3 inhibition in prevention of a broad range of infectious and systemic diseases.

